# Ursolic acid inhibits cell migration and promotes JNK-dependent lysosomal associated cell death in Glioblastoma multiforme cells

**DOI:** 10.1101/2020.03.11.987578

**Authors:** Gillian E. Conway, Deimante Zizyte, Julie Rose Mae Mondala, Zhonglei He, Lorna Lynam, Mathilde Lecourt, Carlos Barcia, Orla Howe, James F Curtin

**Affiliations:** School of Food Science and Environmental Health, Technological University Dublin, Dublin, Ireland; Environmental Sustainability and Health Institute (ESHI) and FOCAS Research Institute, Technological University Dublin, Dublin, Ireland; In-Vitro Toxicology Group, Institute of Life Science, Swansea University Medical School, Swansea University, Singleton Park, Swansea, Wales, United Kingdom; Institut de Neurociències; Department of Biochemistry and Molecular Biology, School of Medicine, Universitat Autònoma de Barcelona, Barcelona, Spain; School of Biological Sciences and Health Sciences, Technological University Dublin, Dublin, Ireland

**Keywords:** Ursolic acid, cancer, therapeutics, cell death, migration, nutraceutical, cell signalling

## Abstract

Ursolic acid (UA) is a bioactive compound which has demonstrated therapeutic efficacy in a variety of cancer cell lines. UA activates various signalling pathways in Glioblastoma multiforme (GBM), however, the relationship between cell death and migration has yet to be elucidated. UA induces a dose dependent cytotoxic response demonstrated by flow cytometry and biochemical cytotoxicity assays. Inhibitor and fluorescent probe studies demonstrated that UA induces a caspase independent, JNK dependent, mechanism of cell death. Migration studies established that UA inhibits GBM cell migration in a time dependent manner that is independent of the JNK signalling pathway. The cytotoxic insult induced by UA resulted in the formation of acidic vesicle organelles (AVOs), speculating activation of autophagy. However, inhibitor and spectrophotometric analysis demonstrated that autophagy was not responsible for the formation of the AVOs and confocal microscopy identified the AVO’s as lysosomes. Further investigation using isosurface visualisation of confocal imaging determined co-localisation of lysosomes with the previously identified acidic vesicles, thus providing evidence that lysosomes are likely to be playing a role in UA induced cell death.

Collectively, our data identifies that UA rapidly induces a lysosomal associated mechanism of cell death in addition to UA acting as an inhibitor of GBM cell migration.

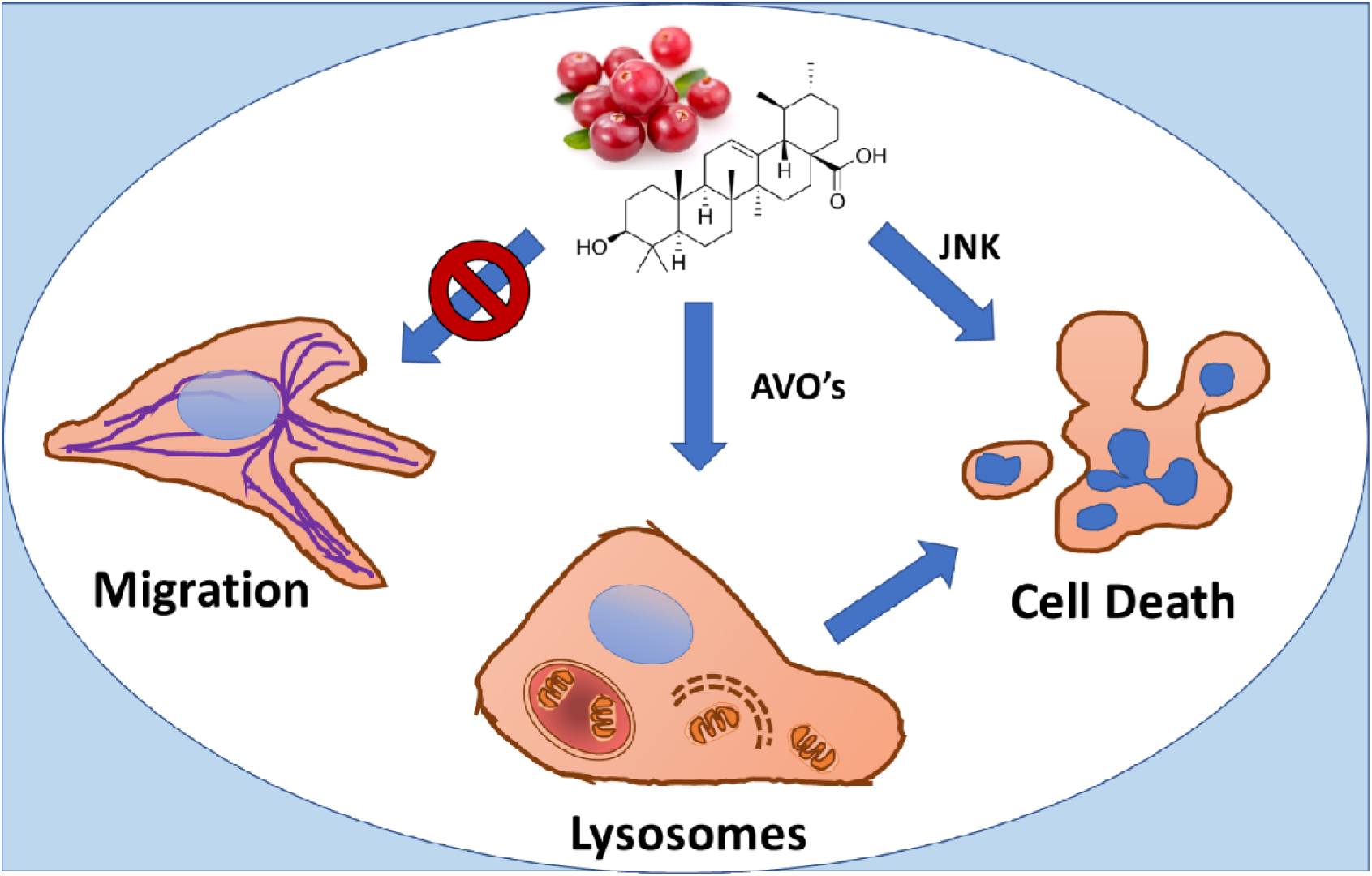

## Introduction

Glioblastoma multiforme (GBM) is a highly malignant grade IV astrocytoma. It is considered to be the most biologically aggressive brain tumour and is associated with a post diagnosis survival rate of only 1 year (Philips et al., 2018). GBMs display attributes of various morphologies such as a heterogeneous mixture of cells that demonstrate varying degrees of cellular and nuclear polymorphism, making it difficult to clinically manage. The current standard of care for GBM is maximal surgical resection followed by radiotherapy and concomitant and adjuvant chemotherapy using Temozolomide (TMZ) (Adamson et al., 2009). The efficacy of chemotherapeutics i.e. Temozolomide (first-line chemotherapeutic for treatment of GBM) and radiotherapy has been limited due to treatment toxicity and therapeutic resistance, and failure to successfully remove all surrounding tumour cells during surgical resection (Candolfi et al., 2007), thus resulting in a high chance of tumour recurrence. Consequently, survival rates have remained relatively stagnant over the past 30 years (Nørøxe et al., 2016). In order to improve clinical outcomes for patients undergoing therapy, there is a need for the development of new therapeutic strategies that investigate alternative compounds that possess cytotoxic activity with the ability to inhibit tissue invasion.

In an effort to overcome these issues, research groups have identified numerous naturally derived bioactive compounds (NDBC) known as phytochemicals that demonstrate anticancer activity when used alone or in combination for the treatment of cancer (Philips et al., 2018). Phytochemicals with anti-cancer properties are being assessed as precursors for new chemotherapeutic agents (Koldaş et al., 2015; Neto, 2007; Øverby et al., 2014; Sasidharan et al., 2011; H. Wang et al., 2012). This is not a new concept, for example, the chemotherapeutic agents topotecan and irinotecan are two derivatives of camptothecin, a bioactive compound isolated from the bark of the *Camptotheca acuminata* tree (Haustedt and Siems, 2015). An international population-based case-control study found that phytochemicals found in yellow/ orange and leafy green vegetables correlate inversely with the incidence of glioma (Terry et al., 2009). Ursolic acid (UA), a ursane-type pentacyclic triterpenoid acid, is a primary component of the thin waxy coating in popular western foods such as cranberries, apples and olives. (Bergamin et al., 2017; Lu et al., 2014; Shanmugam et al., 2013; J. Wang et al., 2012). UA and other triterpenoids are also abundant in Rosemary which has been used in Chinese Traditional Medicine for centuries to treat cancer and inflammatory disorders (Huang et al., 1994). In recent years, UA has been shown both *in vitro* and *in vivo* to demonstrate anti-proliferative and anti-migratory properties, anti-inflammatory, suppress tumour growth and enhance the efficacy and sensitivity of chemotherapeutics in a variety of cancer types including GBM (Navin and Mi Kim, 2016; Woźniak et al., 2015).

UA offers a promising strategy for the treatment of GBM; however, there is a lack of consistency concerning the mechanism of action in the current literature. In GBM models, low concentrations of UA and natural esters of UA inhibited proliferation of U-87 and SF-295 cells (Bonaccorsi et al., 2008; Kondo et al., 2011) and prevented invasion induced by IL1β and TNFα in C6 rat glioma cells (Huang et al., 2009). UA has been shown to induced necrosis in TMZ resistant DBTRG-05MG cells (Lu et al., 2014), cell cycle arrest and autophagy in U-87 cells (Shen et al., 2014), apoptosis in U251 cells (J. Wang et al., 2012), and both anti-proliferative and apoptotic effects in C6 rat glioma cells *in vitro* (Bergamin et al., 2017). An advantage of many naturally derived compounds such as UA is they have the capacity to elicit a biological response while being nontoxic at low doses compared to commonly used chemotherapeutics (Kotecha et al., 2016), making them an ideal candidate for combinational therapy to act as a chemosensitizer. In a recent study, treatment with UA increased the sensitivity to TMZ in TMZ-resistant GBM cells (T98G and LN18) *in vitro* and significantly reduced xenograft *in vivo* both by down inhibition of MGMT (Zhu et al., 2016).

Several key questions have surfaced following observations that UA activates a variety of signalling pathways in GBM. A direct comparison of UA with existing GBM therapies needs to be comprehensively undertaken. The interplay between various anti-tumour signalling pathways, including apoptosis, autophagy and migration is unknown. This study investigates the role of UA induced cell death and its effects on GBM migration, and aims to elucidate the key rate-limiting steps using specific inhibitors.

## Materials & Methods

### Cell culture

Human glioblastoma (U373MG-CD14) cells were obtained from Dr Michael Carty (Trinity College Dublin). U373MG cells, were cultured in DMEM (Sigma-Aldrich, Arklow, Ireland) supplemented with 10% FBS (Sigma-Aldrich, Arklow, Ireland) which were maintained in a humidified incubator containing 5% CO_2_ at 37°C. Media was changed every 2-3 days until 80% confluency was reached. Cells were routinely sub-cultured using a final 1:1 ratio of 0.25% trypsin (Sigma-Aldrich, Arklow, Ireland) and 0.1% EDTA (Sigma-Aldrich, Arklow, Ireland).

### Cytotoxicity

Dose response curves for commonly employed chemotherapeutic drugs used for the treatment of GBM: Temozolomide (TMZ) (Sigma-Aldrich, Arklow, Ireland), Carmustine (BCNU) (Sigma-Aldrich, Arklow, Ireland) and Gefitinib (Insight Biotechnology ltd, Wembley, UK) UA standard (Sigma-Aldrich, Arklow, Ireland) were established. UA standard, TMZ and Gefitinib were dissolved in Dimethyl sulfoxide (DMSO) (Sigma-Aldrich, Arklow, Ireland), and BCNU in sterile H_2_O and stored in −20°C. These stocks were subsequently used to make the working standard solutions in media. The highest concentration of DMSO used was 0.5%. U373MG cells were seeded at a density of 1×10^4^ (24 and 48hr exposure time points) in 96 well plates (Sigma-Aldrich, Arklow, Ireland) with 100µl media per well. In order to establish accurate IC_50_ values for the chemotherapeutic compounds, a lower seeding density of 2.5×10^3^ and a 6-day exposure period was required. Plates were left overnight in the incubator at 37°C with 5% CO_2_ to allow the cells to adhere. Existing media was removed from each well and cells were treated with either a chemotherapeutic drug, UA or solvent control (0.5% DMSO) and incubated for the appropriate time point.

### Cell viability assays

Trypan blue cell exclusion assay was performed as an initial evaluation of cell health following treatment with UA. Cell were trypsinized as described above. A Trypan Blue (Fischer Scientific, Ballycoolin, Ireland) cell suspension was made counted using a haemocytometer as per manufactures instructions.

Cell viability was also measured biochemically using the Alamar Blue assay (Fischer Scientific, Ballycoolin, Ireland). Alamar blue is an oxidation-reduction (redox) fluorogenic indicator of cellular metabolic reduction. After each exposure time point (24hrs or 48hrs) cells were washed once with sterile PBS. A 10% Alamar blue solution in the DMEM was added to each well and incubated at 37°C for 2.5 hours. Fluorescence was read at an excitation wavelength of 530nm and an emission wavelength of 595nm using the Victor 3V 1420 (Perkin Elmer) multi-plate reader.

### Flow Cytometry

#### JC-1 mitochondrial membrane potential assay

As described above cells were plated and exposed to varying concentrations of UA for 48 hours. Cells were then harvested and stained with 10µg/ml JC-1 dye, a mitochondrial membrane potential probe (Biosciences, Dublin, Ireland) (Galluzzi *et al*, 2007), at room temperature for 10 minutes and analysed by flow cytometry (BD Accuri C6). JC-1 was excited using the argon laser at a wavelength of 488 nm. Fluorescence was measured using the FL1 (530 nm) and FL2 (585 nm) channels with emission spectral overlap compensation (7.5% FL1/FL2 and 15% FL2/ FL1).

#### Acridine orange (AO)

Cells were harvested and stained with 1µg/ml AO and incubated at 37°C for 20 minutes. Cells were then washed twice with sterile PBS and analysed by flow cytometry (BD Accuri C6) with an excitation of 475nm and emission 590nm. Detection of AVO’s was achieved using the FL1 (green) vs FL2 (orange) channels and compensation was set at approximately at 7% removing FL2 signal from FL1 and approximately 16% removing FL1 signal from FL2 for each plot.

### Inhibitor studies

U373MG Cells were plated in 96 well plates as described above and left to adhere overnight. Cells were pre-treated for 1 hour with either zVAD-FMK; caspase inhibitor (BD Bioscience, Oxford, England), SP600125; JNK inhibitor, SB203580; p38 MAP Kinase Inhibitor (Sigma Aldrich, Arklow, Ireland), U0126; MEK1 and MEK2 Kinase Inhibitor (Sigma Aldrich, Arklow, Ireland), chloroquine or 3-methyladenine (3-MA), autophagy inhibitors (Sigma Aldrich, Arklow, Ireland) after which the UA standard (IC50) was added to each well. Cell viability was assessed 48 hours later using Alamar Blue cell viability assay.

### Confocal Microscopy

U373MG cells were plated in 35mm glass bottom dishes (MatTek Corporation, USA) at 1×10^5^ cells per ml and incubated for 24 hrs. Cells were treated with 20µM UA for 48hrs. After treatment, cells were loaded with AO (1µg/ml for 15minutes at 37°C) and LysoTracker Deep Red (Thermo Fisher Scientific) (50nM for 30 minutes at 37°C). Images were captured on a Zeiss 510 LZSM confocal inverted microscope. The corresponding filter settings were as follows: AO, excitation 477 nm, emission 585-615 nm; LysoTracker Deep Red, excitation 633 nm, emission 649-799 nm. All images were taken using live cells. The total integrated density of fluorescence of each cell was quantified using ImageJ (v1.49, NIH) software. The quantified integrated density equals to the sum of the pixel values in the selected fluorescent area.

### Cell Migration Assay

The scratch assay allows for the preliminary examination of the effects of TMZ, Gefitinib, BCNU, and UA on the migration of U373MG cells. Cells were treated with each compound, at a subtoxic concentration; to prevent cytotoxic responses but potentially inhibit migration. U373MG cells were seeded at 0.9 × 10^6^ cells in individual 35mm dishes and incubated for 24hrs. A scratch was performed in each dish prior to treatment using a 200µl sterile pipette tip. Utilising the data observed from the UA dose response curve a sub-toxic concentration (12μM) was chosen. Cells were treated with either media alone, DMSO (0.1%), or 12.5μM UA, each scratch was examined under a light microscope and images were taken using the (Nikon Eclipse 700). Multiple images were taken for each time point and the average size of scratch for that time point was obtained. Image analysis was performed using image processing and analysis software, Image J (Schneider et al., 2012).

### Isosurface rendering

Representative confocal Z-scans of U373MG cells were processed for a three-dimensional reconstruction and visualization of volumetric co-localization of the acidic vesicles (ACRID) and lysosomes (LYSO). Fluorescent ACRID-LYSO colocalizing voxels were detected by a computerized software using Coloc module (Imaris Bitplane, South Windsor, USA), then a new channel with these voxels was extracted. Then, this colocalizing voxels channel was used to generate a three-dimensional isosurface employing Surface module (Imaris BitPlane, South Windsor, USA) with the appropriate threshold and resolution. Similarly, ACRID and LYSO channels were also used to generate three-dimensional renderings as isosurfaces. Merging of the volumetric isosurfaces was set to illustrate the degree of colocalizing objects in control and treated glioma cells. Adequate shadowing and angle of three-dimensional rotation was applied to show the relevant cellular structures.

### Statistical Analysis

All experiments were performed in triplicate, independently of each other with a minimum of five replicates per experiment. Data shown is pooled and presented as mean ± SEM (n= total number of replicates) unless stated otherwise. Statistical analysis was performed using Prism 5, GraphPad Software, Inc. (USA). Unless otherwise indicated differences were considered significant with a *P value < 0.05.

## Results

### UA induced cytotoxicity results in rapid mitochondrial membrane depolarisation

UA was found to induce rapid cytotoxicity in U373MG GBM cells within 24hrs of treatment. The IC_50_ of 18 µM (95% confidence range: 17.09 µM to 19.46 µM) is consistent that observed in other GBM cell lines such as U251, DBTRG-05MG and C6. The rapid onset of cell death was accompanied by a characteristic, steep Hill slope, with no significant difference in IC_50_ values determined when cells were treated with UA for 24 hrs, 48 hrs or 6 days (R^2^ > 0.9736 > 0.05) (Figure 1a). To help better understand UA induced cytotoxicity, three concentrations (1.5 µM UA (Low), 20 µM (IC50) or media only (Control)) were chosen for evaluation using the Trypan blue cell exclusion assay 48 hrs post treatment. As expected, there is a significant (P<0.0001) increase in the number of dead cells following treatment with UA IC_50_ in comparison with the untreated control (Figure 1b, left panel). Interestingly, when treated with low doses of UA (1.5μM), there is significant increase (P<0.001) in live cell number compared to cell treated with the UA IC_50_(Figure 1b, right panel). It is hypothesized therefore that low doses of UA may promote cell proliferation, due to mild oxidative stress induced by UA (Gupta et al., 2012; Shen et al., 2008). Mitochondrial membrane potential (ΔΨm) is an important factor of mitochondrial function and can be an indicator of early intrinsic apoptosis. Collapse of the ΔΨm results in the release of cytochrome C into the cytosol and thus leading to cell death (Salido et al., 2007). Mitochondrial involvement in cell death was explored using the voltage sensitive fluorescent probe JC-1. Loss in ΔΨm was observed following treatment with UA (P < 0.05) in a dose dependent manner (Figure 1c), with a significant loss in ΔΨm with 25 µM compared to the untreated control. This data correlated with the loss of respiration function measured using Alamar blue and indicated that depolarisation of mitochondria is an early feature of cell death induced by UA.

**Figure 1.**
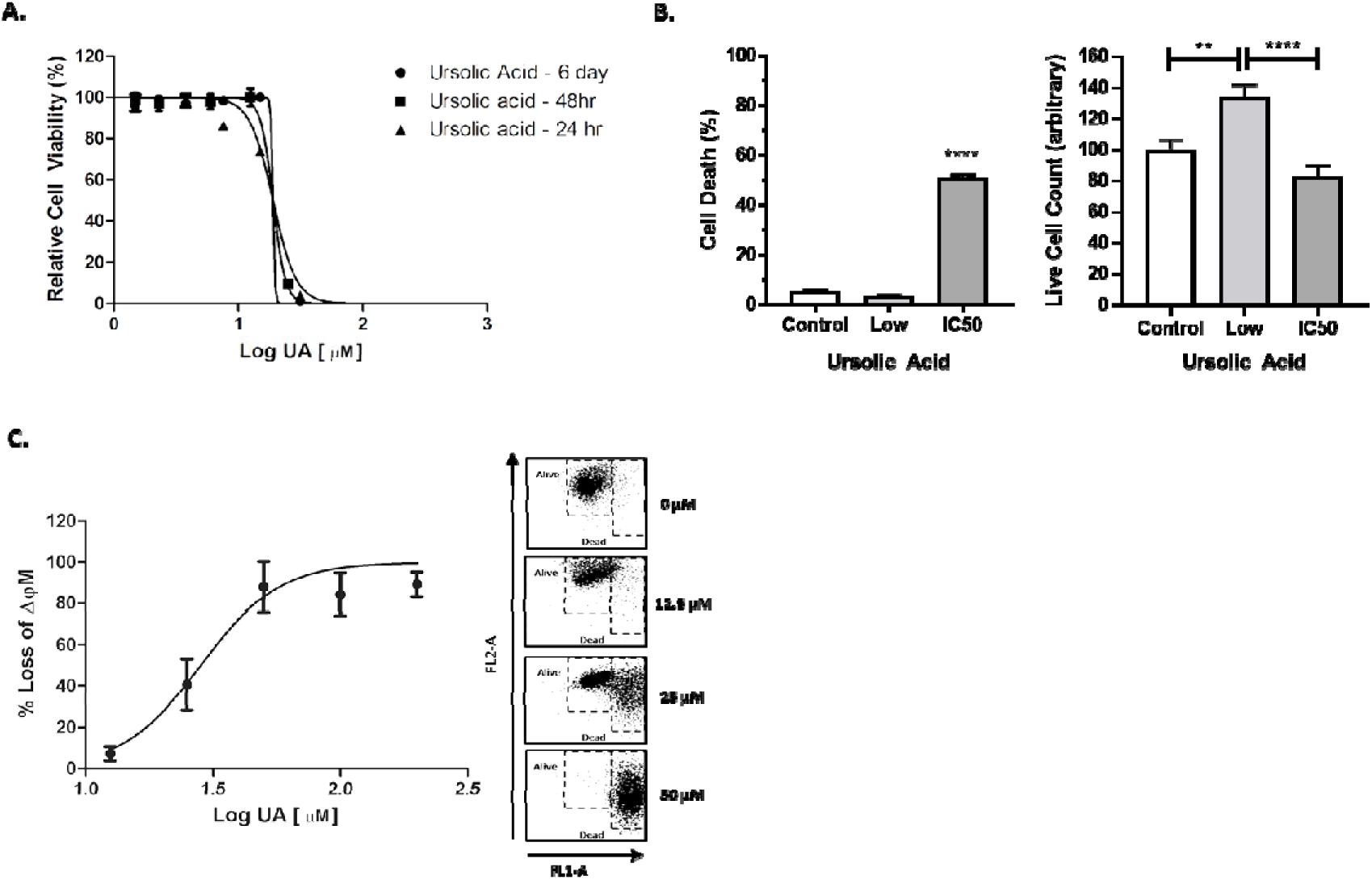
UA induces mitochondrial membrane depolarisation. (**A**) U373MG cells were exposed to increasing concentrations of UA. Cell viability was assessed using Alamar blue cell viability assay at 24, 48 and 6 days (n=3). (**B**) U373MG cells were treated with 1.5 µM UA (Low), 20 µM (IC50) or media only (Control) for 48 hrs. Cell health was measured using the trypan blue cell exclusion assay (n=3) **(C**) U373MG cells were exposed to increasing concentrations of UA (standard). After a 48hrs incubation period, cells were loaded with 10µg/ml JC-1 dye and analysed by flow cytometry. Data shown depicts cell death measured by quantitative shifts in the ΔΨm (red to green) fluorescence intensity ratio with increasing concentrations of UA (n=3). Data shown was normalised to the untreated control and are shown as the % mean ± S.E.M. Statistical analysis was carried out using non-linear regression.

### UA demonstrates enhanced cytotoxicity compared to conventional chemotherapeutic drugs

The cytotoxicity of UA was also compared to the standard chemotherapeutic drugs used for the treatment of GBM (TMZ) and for recurrent disease (Gefitinib and BCNU) (Gallego, 2015). Little cytotoxicity was observed 48 hrs after treatment with Gefitinib, TMZ or BCNU, which prevented IC_50_ values being accurately calculated. In contrast, UA demonstrated a significant reduction in cell viability 48 hrsafter treatment (Figure 2a), with an IC_50_ value of 22µM, similar to that observed in figure 1a. Cytotoxicity was observed 6-days post treatment using TMZ, BCNU and Gefitinib, (IC_50_ values were 28 µM, 79 µM and 16 µM, respectively) in comparison with UA (19 µM) (Figure 2b). Interestingly, UA remains stable over the 6 days period with only a small reduction in the IC_50_ value. As individual agents, UA demonstrated greater cytotoxicity over shorter period and at significantly lower concentrations then that used for TMZ. To date there has been only one published article that has investigated the synergistic effects of UA with TMZ. The combined treatment of TMZ and UA synergistically enhanced the cytotoxicity and senescence in TMZ-resistant GBM cells. The study also demonstrated a reduction in tumour volume by depletion of MGMT *in vivo*, demonstrating that UA down regulates MGMT status in GBM cells (Zhu et al., 2016). As demonstrated below in Figure 2c, while there is a significant difference between with and without TMZ, no additive or synergistic effect was observed between low doses of TMZ and UA. We postulated that as U373MG are TMZ sensitive, and therefore UA did not have a demonstrable effect on MGMT.

**Figure 2.**
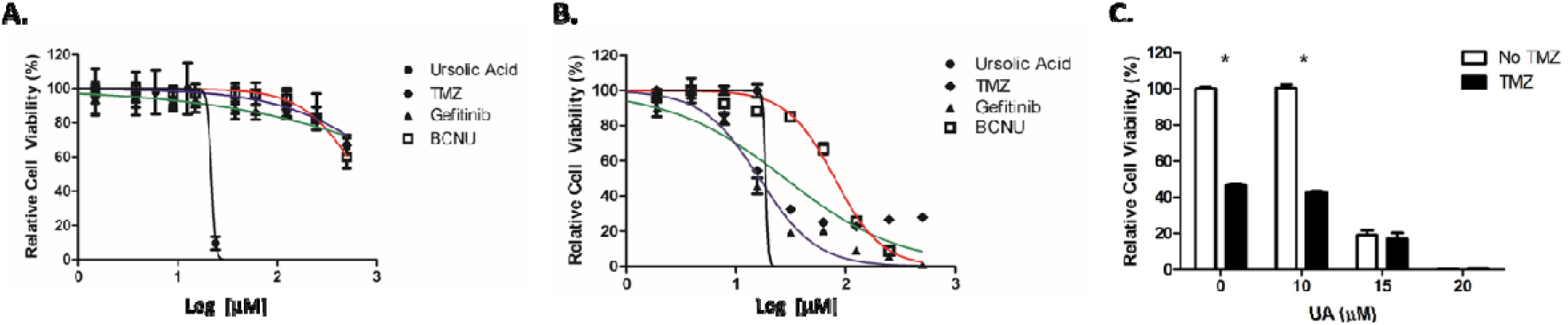
UA exhibits increased cytotoxicity over conventional chemotherapeutics. (**A, B**) U373MG cells were treated with increasing concentrations of UA (0-200μM), TMZ, BCNU, or Gefitinib (0-500μM) for 48 hours (A) or 6 days (B) and analysed using Alamar blue cell viability assay. Statistical analysis was carried out using non-linear regression analysis and Two-way ANOVA with Bonferroni post-tests, (n=3) (*P<0.001). (**C**) U373MG cells were treated with 0-20μM UA after which 15μM TMZ was added to the wells. Cells were then incubated for 6 days and analysed by Alamar blue. Cells were also treated with a vehicle control of 0.2% DMSO. No deleterious effects were observed. Data shown was normalised to the untreated control and are shown as the % mean ± S.E.M. Statistical analysis was carried out using Two-Way ANOVA with Bonferroni post-test (*P<0.001) (n=3).

### UA induces caspase-independent, JNK-dependent cell death

The rapid onset of cell death and a steep Hill slope relative to TMZ, BCNU and Gefitinib indicated that an alternative mechanism of cell death may be activated following treatment with UA. Several competing hypotheses have been put forward regarding the signalling pathways responsible for UA induced cytotoxicity, with both caspases and kinases being implicated (Gao et al., 2012; Kassi et al., 2007; J. Wang et al., 2012; Xavier et al., 2013; Zhang et al., 2010). As mentioned above the Hill slope for UA is high with small changes in concentration around the IC_50_ value leading to large changes in viability. Therefore, three concentrations of UA were tested in order to capture low, moderate and high degrees of cytotoxicity. The JNK specific inhibitor SP600125 alleviated cytotoxicity in U373MG cells induced by moderate concentrations of UA (20µM) (p<0.001) with a resultant significant increase in the IC_50_ value (p<0.05) (Figure 3a). There was no evidence of any inhibitory effects by the broad-spectrum caspase inhibitor zVAD-fmk as seen in Figure 3b, indicating that caspase enzymatic activity may not be playing a role in UA-induced cytotoxicity in U373MG cells. Having identified that JNK plays a role in UA induced cell death in GBM cells; it was hypothesized whether stress-activated p38 MAPK signalling pathways were involved. It was previously identified that UA-induced apoptosis is regulated by p38 and JNK and MAPK signalling pathways in both liver and gastric cancer cells (Chuang et al., 2016; E.-S. Kim and Moon, 2015). However, as demonstrated in Figure 3c, there was no evidence of any inhibitory effects by the p38 kinase inhibitor SB203580. In agreement with previous reports, Figure 3d, demonstrated that JNK was involved in TMZ-induced cytotoxicity (p<0.05), in contrast to this, no inhibition was observed when TMZ was combined with Zvad-fmk. These results suggest that UA induces a rapid, Caspase- and p38-MAPK independent, JNK dependent mechanism of cell death in U373MG GBM cells that involves mitochondrial membrane depolarisation.

**Figure 3.**
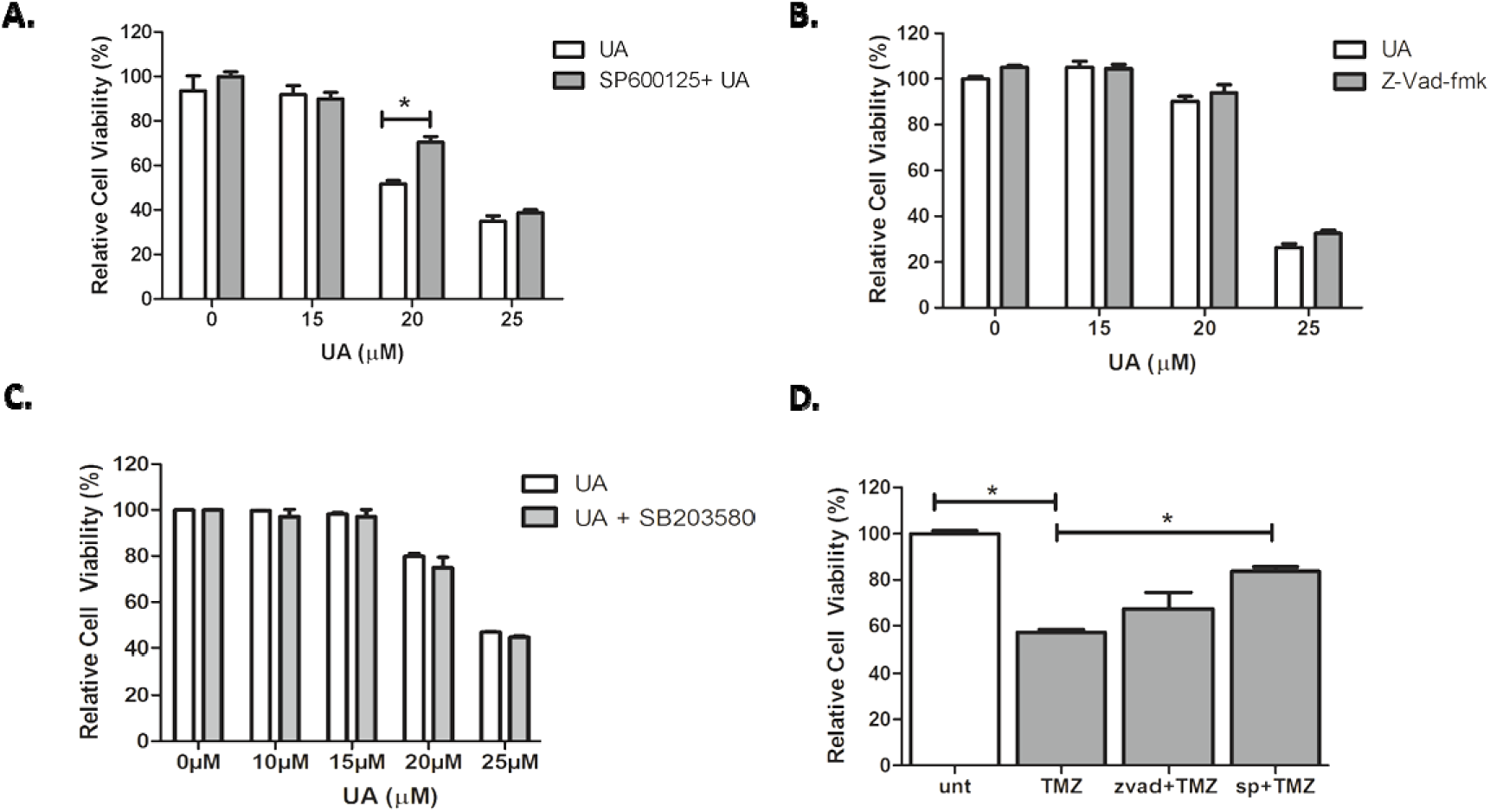
UA induces Caspase-independent, JNK-dependent cell death. (**A**) Cells were pre-loaded 12.5µM SP600125 JNK inhibitior for 1 hr, increasing concentrations of UA were added in the presence of 12.5µM SP600125 and incubated for 48hrs. Cells were then analysed using Alamar blue cell viability assay. Statistical analysis was carried out using Two-Way ANOVA with Bonferroni post test. (**B**) U373MG cells were pre-treated with 50µM zVAD-FMK for 1hr prior to UA treatment. Cells were then incubated for 48hrs and analysed by Alamar blue. Statistical analysis was carried out using Two-Way ANOVA with Bonferroni’s post test. (**C**) Cells were pre-loaded 10µM SB203580 p38-MAPK inihibitor for 1 hr, increasing concentrations of UA were added and incubated for 48 hrs. Cells were then analylsed using Alamar blue cell viability assay. Statistical analysis was carried out using Two-Way ANOVA with Bonferroni post test (**D**) Cells were pre-treated with either zVAD-fmk (50μM) or SP600125 (12.5μM) for 1 hrprior to addition of TMZ (26.5μM) for 6 days. Cell viability was analysed by Alamar blue. Statistical analysis was carried out using One-Way ANOVA with Tukey’s post-test (p<0.05). All experiments were normalised to untreated control and expressed as a % of the SEM (n=3).

### UA inhibits JNK-dependent GBM cell migration

One of the most important hallmarks of GBM is the invasive behaviour and ability to migrate into surrounding healthy brain tissue. Despite surgical resection, statistics show that the tumour recurs within 1cm of the surgical site (Demuth and Berens, 2004). Sub-toxic doses of UA (10-20 μM) abrogated IL1β- and TNFα-induced GBM cell migration in C6 rat glioma cells by inhibiting PKCζ activation and reducing MMP-9 expression (Huang et al., 2009). Our data demonstrates that JNK activation promotes caspase-independent UA cytotoxicity and interestingly, JNK has also previously been implicated in promoting cell migration (Zhou et al., 2012). We therefore hypothesised that sub-toxic concentrations of UA, following activation of JNK signalling, might inadvertently promote or inhibit GBM cell migration.

Treatment of U373MG cells with SP600125 combined with UA reduced the rate of migration compared with untreated controls (p < 0.05, Figure 4a). Migration was reduced to a lesser extent when cells were treated with SP600125 alone, which correlates with results observed by Yeh *et al* (C. T. Yeh et al., 2010). As previously identified above (Figure 1b), no toxicity was observed with 12.5μM UA after 24 hrs, however, this may have an effect on migration signalling. Our data demonstrates that 12.5µM alone significantly reduced the rate of migration compared with untreated control (p<0.05, Figure 4a). No cumulative effect was observed when UA and SP600125 were combined (p>0.05), indicating a role for UA as an inhibitor of GBM cell migration.

**Figure 4.**
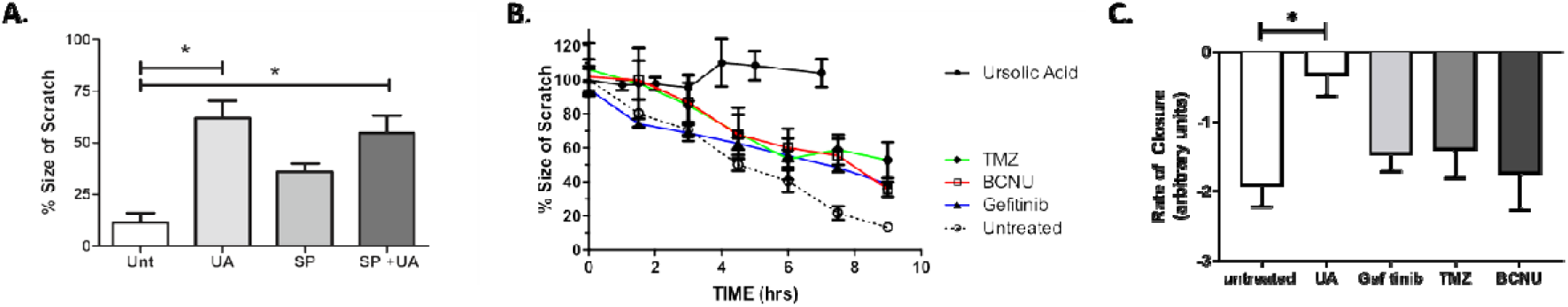
UA inhibits JNK-dependent GBM cell migration. (A) The effects of JNK inhibition using SP600125 on scratch closure when treated with UA. Cells were treated with either, 12.5μM UA, SP600125 12.5μM alone and UA/SP combined. Images were taken at the time of the scratch and 7 hrs later. Image analysis was performed using ImageJ software. Statistical analysis was carried out using One-Way ANOVA with Tukey’s multiple comparison post test (*P<0.05) (n=3). (**B&C**) U373MG cells were either left untreated, or treated with TMZ (100μM), Gefitinib (25μM), BCNU (50μM), or UA (12μM). Time dependent closure of the scratch was then analysed over a 9-hrs period. Image analysis was performed using image J Software. Statistical analysis was carried out using linear regression analysis and Two-Way ANOVA with Bonferroni post-tests. The rate of closure was determined by One-Way ANOVA with Tukey’s test for multiple comparisons (n=3).

Next, a comparison of UA and currently used chemotherapeutic drugs to inhibit migration was investigated using the scratch assay. The Pearson’s correlation coefficient was calculated over the course of 9 hrs for untreated U373MG cells and for cells treated with sub-toxic concentrations of TMZ, BCNU, Gefitinib and UA. The correlation coefficient was very strong and negative (i.e. r > - 0.95) for both untreated and cells treated with TMZ, BCNU and Gefitinib, thus indicating treatment with these chemotherapeutic agents did not affect cell migration. The correlation coefficient for cells treated with UA was positive, indicating that UA had completely blocked migration (see Table 1).

**Table 1.** Investigation of the effect of UA on U373MG cell migration. Correlation Coefficients for cells treated with UA for up to 9 hrs.

| Treatment | Correlation Coefficient (r) | P value (two-tailed) |
| --- | --- | --- |
| Untreated | -0.9635 | 0.0020 |
| Gefitinib | -0.9422 | 0.0049 |
| TMZ | -0.9676 | 0.0016 |
| BCNU | -0.9699 | 0.0013 |
| UA | 0.6627 | 0.1515 |

Statistical analysis confirmed that a significant level of migration was observed for untreated cells and for cells treated with TMZ, Gefitinib and BCNU (p<0.01, Figure 4b), whereas no significant evidence of migration was observed for cells treated with UA (P>0.05). To further examine the changes in migration following treatment with UA, the rate of scratch closure was then calculated. Figure 4c and supplemental movie of scratch closure demonstrates that treatment of GBM cells with UA significantly (p< 0.001) reduces the rate of closure of the scratch when compared to untreated control. No significance was observed for any chemotherapeutic agents when comparted to untreated cells. These data support our findings that UA is a potent *in vitro* inhibitor of GBM cell migration.

### UA triggers the formation of lysosomes

Autophagy is an essential cytoprotective response to pathological stressors, involving the phosphatidylinositol 3-kinase PI3K/AKT/mTOR signalling pathway and also plays a role tumorigenesis and invasion of tumour cells (Murrow and Debnath, 2013). It was previously identified that UA stimulated autophagy in response to ROS-mediated ER stress in U-87MG cells (Shen et al., 2014). Having identified that UA induces cytotoxicity response that is independent of caspases but dependent on JNK signalling, it was postulated whether autophagy was activated in response to UA treatment.

Following treatment with UA, U373MG cells were stained with Acridine orange which is used for the detection of acidic vesicle organelles (AVO’s). AVO’s are deemed a significant morphological characteristic of the autophagic process, (Paglin et al., 2001; Shin et al., 2012). As seen in Figure 5a, there is a significant increase (P<0.005) in the formation of AVO’s following UA treatment compared with untreated cells. This data suggests that the autophagy pathway may be activated during UA-induced cell death. However, as demonstrated below, upon further investigation no evidence of autophagy activation was identified. 3-MA is an inhibitor of autophagosome formation through Class III PI3K activity which is critical for autophagy activation. As demonstrated in Figure 5b, no evidence of inhibition was observed. Having previously identified that JNK is playing a role in UA-mediated cell death, it was postulated whether other MAP kinases (MEK1 and MEK2) might be activated, as they have previously been shown to facilitate induction of autophagy (Li et al., 2017). No inhibition of cytotoxicity was observed following addition of U0126 inhibitor (Figure 5c), indicating that UA induced cytotoxicity is independent of ERK activation. Autophagic flux is used as a measure of autophagic degradation, thus, is associated with late stage autophagy. Cells were treated with chloroquine a well-known inhibitor which works by destroying autophagosome and lysosome fusion (Mauthe et al., 2018). Chloroquine was unable to prevent cytotoxicity induced by UA (Figure 5d). Together these results demonstrate that UA-induced AVO formation reminiscent of autophagosomes but cytotoxicity was not responsive to common modulators of autophagy.

**Figure 5.**
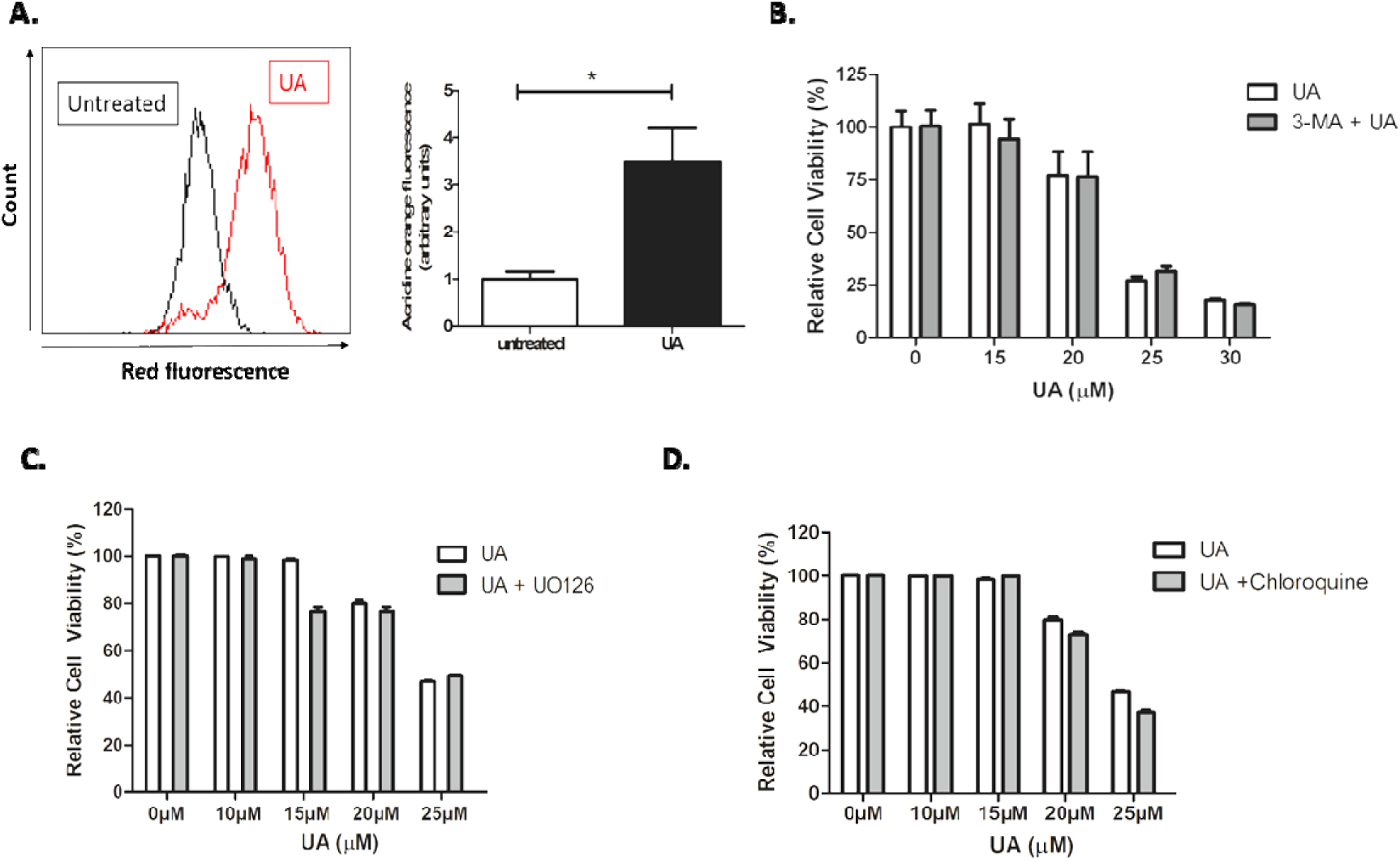
UA treatment does not initiate an autophagic response. **(A**) U373MG cells were exposed to 22µM UA. After a 48 hrs incubation period, cells were loaded with 1µg/ml acridine orange fluorescent probe and analysed by flow cytometry. Data shown depicts a significant increase in the presence of AVO’s measured by quantitative shifts in the FL-2 channel (red fluorescence) intensity ratio following treatment with UA. Data shown was normalised to the untreated control and represented as arbitrary units ± S.E.M (n=3). Quantification of mean fluorescence index and statistical analysis using non-parametric t-test (*P<0.05). (**B-D**) U373MG cells were pre-treated with 5mM 3-MA, 10µM U0126, or 25µM Chloroquine for 1 hr prior to addition of UA. Cells were then incubated for 48 hrs and analysed by Alamar blue. Data shown was normalised to the untreated control and are shown as the % mean ± S.E.M (n=3). Statistical analysis was carried out using Two-Way ANOVA with Bonferroni post-test.

It was postulated whether the acidic vesicles identified in Figure 5a maybe an alternative acidic vesicle such as lysosomes. Lysosomes are acidic vesicles that contain hydrolytic enzymes which function to recycle damaged organelles. Permeabilisation of the lysosomes following damage to the lysosomal membrane results in the release of the hydrolytic enzymes and cathepsins, which then results in a process called lysosomal dependent cell death (LDCD). Figure 6a demonstrates a significant increase of acidic vesicles following treatment with UA. However interestingly when stained with lysosomal tracker deep red (Figure 6b) there is also a significant increase in fluorescence intensity following UA treatment (p<0.0001). This data provides evidence that the vesicles detected using AO are lysosomes. To confirm the presence of lysosomes, 3D isosurface rendered z-stack confocal images were generated from cells treated with both AO and LysoTracker Deep Red. Figure 6c demonstrates co-localisation, as depicted in blue in the merged panel. As previously described, AO also binds with single-stranded nucleic acids and emits orange fluorescents which can be seen in both the merged panel of both the untreated control and UA treated cells (Conway et al., 2019).

**Figure 6.**
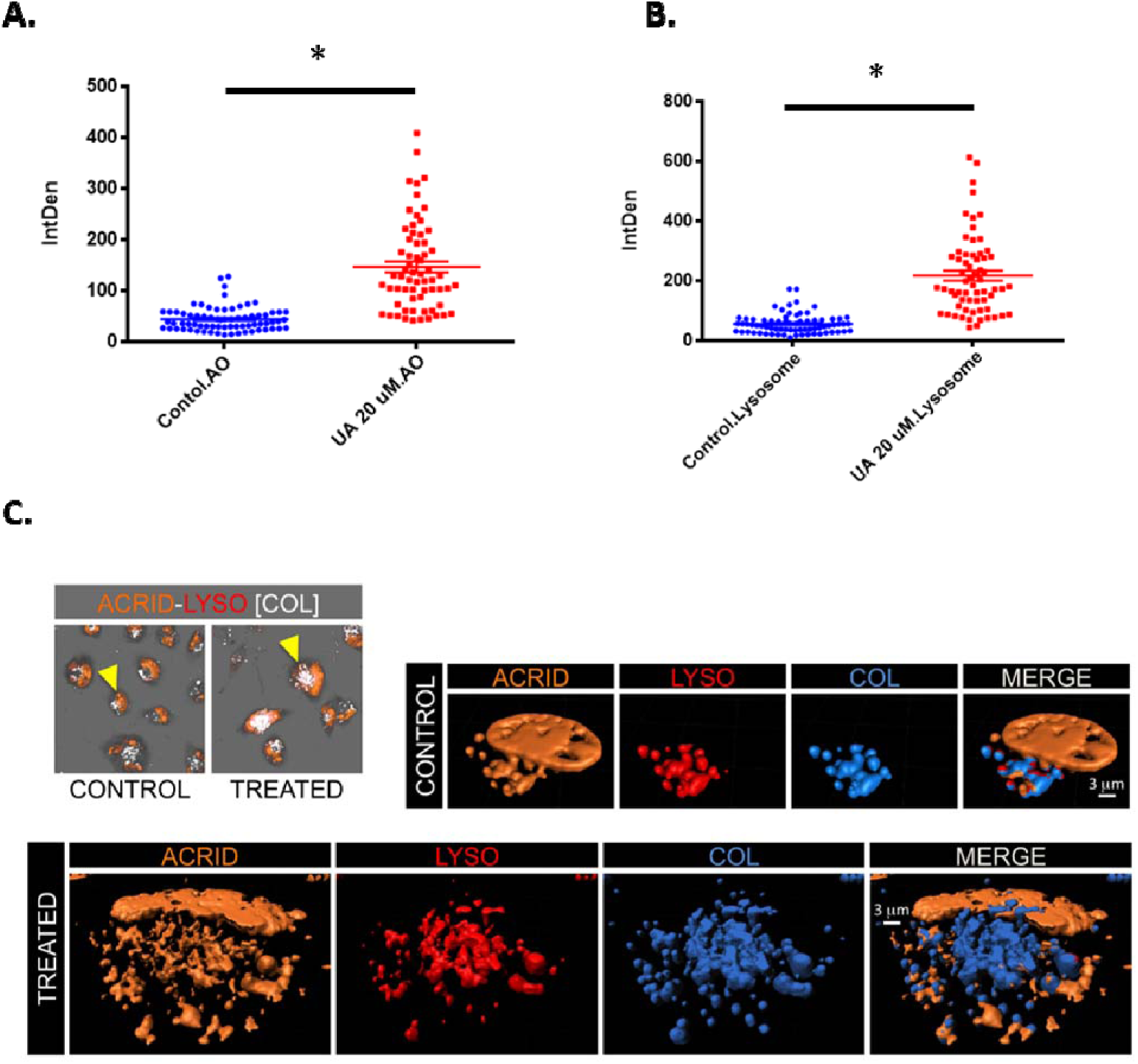
UA triggers lysosome accumulation. **(A)** U373MG cells were exposed to 20µM UA. After a 48 hrs incubation period, cells were loaded with acridine orange (1µg/ml for 15 minutes at 37°C) and (**B**) 50nM lysotracker (50nM for 30 minutes at 37°C) and analysed by confocal microscopy. The fluorescence intensity of AO green and orange channel, and LysoTracker Deep Red was quantified using ImageJ software and compared to the untreated control. Statistical analysis was carried out using an unpaired t-test (*p<0.0001). (**C**) Images showing co-localisation analysis of Lysotracker Deep Red (LYSO) and AO (ACRID) channels as white voxels (COL) for both control and UA treated cells. 3D visualization of control and treated cells demonstrates the co-localisation of orange (acridine orange) and red (Lysotracker Deep Red) indicated with a blue isosurface (COL) toguether with rendered isosurface of the AO (ACRID) and Lysotracker (LYSO) in separated channels or merged images (MERGE)..

## Discussion

An estimated 25% of chemotherapeutic drugs used during the last 20 years are directly derived from plants, while another 25% are chemically altered from natural products. It is noteworthy, however, that only 5-15% of the approximately 250,000 higher plants have ever been investigated for bioactive compounds (Amin et al., 2009). UA, found in the leaves and fruits of many plants, induces cytotoxicity in cancer cells while remaining nontoxic to normal cells (Shanmugam et al., 2013; Weng et al., 2014). Derivatives of UA have also demonstrated increased antitumor efficacy *in vivo* (Wang et al., 2011; Yoon et al., 2016; Zhang et al., 2015). This paper demonstrates that UA induces JNK-dependent and caspase independent cytotoxicity in a human GBM model. Other groups have reported various cytotoxic and pro-survival signalling pathways activated by UA in GBM cancer cell lines, including apoptosis, autophagy, senescence and necrosis. This variability in responses observed in GBM appears partly dependent on the treatment regimen, relative resistance to TMZ and the GBM model used. Lower doses of UA were generally anti-proliferative in GBM cells without directly inducing cytotoxicity (Bonaccorsi et al., 2008; Kondo et al., 2011). Cytotoxicity observed at higher doses of UA does not always carry the hallmark indicators of apoptosis. Biochemical indicators of apoptosis, including caspase activation, phosphatidylserine exposure, and DNA fragmentation were evident when TMZ-sensitive GBM cells such as C6 cells (Bergamin et al., 2017) and U251 cells (J. Wang et al., 2012) were treated with UA *in vitro*. However, treatment of TMZ-resistant GBM cells with UA led to cytotoxicity associated with caspase-independent signalling, such as necrosis in DBTRG-05MG (Lu et al., 2014), LN18 and T98G cells (Zhu et al., 2016). Interestingly, in TMZ-sensitive U87MG cells, autophagy via ROS-induced ER stress was reported in parallel with cytotoxicity (Shen et al., 2014). Meanwhile, Cho and colleagues previously demonstrated that Asiatic acid, a triterpenoid with similar structure to UA, appeared to induce features of both apoptotic and necrotic cell death in U87MG cells, including dose-dependent Caspase 3 and 9 activation, phosphatidylserine flipping, mitochondrial membrane depolarisation and LDH release (Cho et al., 2006).

TMZ resistance is believed to be conferred by MGMT expression in GBM cells and correlates with poor prognosis and U373MG cells are considered TMZ-sensitive (Hermisson et al., 2006). Our findings that UA induces JNK-dependent, caspase independent cytotoxicity in U373MG cells differs to that described in other TMZ-sensitive cell lines. UA was shown to reduce the expression of MGMT in MGMT-expressing cells and induce synergistic cytotoxicity in TMZ-resistant GBM cell lines T98G, LN18 and LN229 when co-incubated with TMZ. Moreover, the rate of tumour growth was reduced when mice challenged with flank LN18 xenograft tumours were treated with UA and TMZ (Zhu et al., 2016). We did not observe any additive or synergistic effect when U373MG cells were treated with a combination of TMZ and UA. This may be due to the steep Hill slope observed when U373MG cells were treated with UA, compared to T98G, LN18 and LN229 cells that display significant resistance to UA and TMZ, or perhaps MGMT expression in U373MG cells is regulated through a different, UA-insensitive signal transduction pathway.

Studies to date using GBM cells suggest that caspase involvement in UA-induced cell death appears to be associated with TMZ-sensitivity of the particular cell lines used and caspase independent cell death observed was described as necrosis. Our data suggests, that caspases are not required for UA-induced cytotoxicity in U373MG cells. Instead, we found that JNK regulates cytotoxicity in GBM cells. No hallmarks of necrosis or necroptosis were observed, such as cell swelling and lysis, suggesting that caspase-independent apoptosis is induced by JNK in U373MG cells in response to UA. JNK-dependent apoptosis involving caspase activation has previously been reported for various human cancer cell lines treated with UA including pancreatic (Liang, 2012), bladder (Zheng et al., 2013) and prostate (Zhang et al., 2010). We found that UA induced JNK-dependent, caspase-independent cell death which correlates with findings observed in human HCT15 colorectal cancer cells (Xavier et al., 2013). Interestingly, we found that SP600125 could significant lower cell death at a concentration close to the IC_50_ of UA (i.e. 20µM), but not at higher doses. Our observations correlate with those observed by Zhang *et al* in prostate cancer cells (Zhang et al., 2010) and may indicate that other cytotoxic signalling pathways can be activated by UA depending on the cell line.

JNK activation has also been implicated during activation of autophagy by UA. Xavier *et al* reported that UA increased expression of the autophagic indicators LC3-II and P62 in colorectal carcinoma cells which could be inhibited using SP600125 (Xavier et al., 2013). Similar findings of ER stress-induced JNK activation and autophagy were reported in U-87MG cells (Shen et al., 2014). In both studies, autophagy was associated with cell death, as was also the case in a study using the human cervical carcinoma cell line TD1, where authors observed UA-induced cell death without a DNA fragmentation, caspase activation or phosphatidylserine flipping. UA-induced LC3-II conversion, autophagosome formation, and inhibition of UA-induced autophagy using Wortmannin or ATG5 siRNA led to a significant improvement in survival. The authors conclude that autophagy is involved in promoting UA cytotoxicity (Leng et al., 2013). The role of autophagy in the cell death process is not fully understood. The term ‘autophagic cell death’ was widely used in the literature, based on the observation that autophagy is commonly associated with cell stress events, and that instances of cell death accompanied by a massive cytoplasmic vacuolization lead to the conclusion that autophagy is an enabler of cell death in cells (Galluzzi et al., 2012). More recently, however, it was noted that inhibition of the processes that are essential for cell death often do not result in inhibition of the cells demise but more often result in alternative biochemical pathways inducing cell death via a different mechanism (Galluzzi et al., 2015).

We aspired to clarify the role of UA in our GBM model and identify the vesicles that were observed following UA exposure. It is important to note that acridine orange cannot be used as an identifier of autophagic vesicles but more as an indicator. Acridine orange crosses into acidic compartments such as AVO’s and lysosomes and becomes protonated. Therefore, it cannot be ruled out that the observed shift in fluorescence is from the uptake of acridine orange by lysosomes neither acidic vesicle organelles that are associated with autophagy. Our data demonstrated no inhibition of autophagy with the PI3 Kinase inhibitor 3-MA. Leng *et al* observed a cytoprotective effect using Wortmannin (a PI3K inhibitor) and ATG5 siRNA in combination with UA. However, they also observed no cytoprotective effect using PI3K inhibitors 3-MA, despite observing inhibition of LC3-II and accumulation with LY294002. Our observation that inhibition of PI3-K using 3-MA did not alleviate the cytotoxic effects of UA correlates with Leng *et al* (Leng et al., 2013). To further confirm that autophagy was not playing a role, our data demonstrates no inhibitory effects of chloroquine on UA treated cells. Chloroquine is lysosomotropic agent that prevents endosomal acidification and therefore inhibits autophagy as it raises the lysosomal pH, leading to inhibition of both fusion of autophagosome with lysosome and lysosomal protein degradation. It was then postulated that the observed influx in acidic vesicle organelles were in fact lysosomes, and thus providing a rationale why no inhibition was observed.

UA has also been implicated in regulating cell migration and invasion in gastric cancer cells, breast and lung cancer cells (Huang et al., 2011; E. Kim and Moon, 2015; C.-T. Yeh et al., 2010). Another compound ‘oleanolic acid’, also a pentacyclic triterpenoid, is structurally similar to UA differing only by the position of one methyl group on the ring E (Ovesná et al., 2006). It has also demonstrated anti-migratory and invasion in two GBM cell lines, U251MG and U87MG, by inactivating the MAPK/ERK signalling pathway (Guo et al., 2013). We observed that UA was a potent inhibitor of U373MG cell migration. UA has also been shown to inhibit chemotaxis in polymorphonuclear leukocyte (PMN) (Mawa et al., 2016) and inhibit migration of breast (C.-T. Yeh et al., 2010) and lung cancer cells (Huang et al., 2011). Investigation in human breast cancer cells found that migration was affected through JNK, Akt and mTOR signalling (C.-T. Yeh et al., 2010). A possible link between JNK, migration and autophagy was tantalising in our case. However, while we observed that inhibition of JNK reduced U373MG cell migration, we did not see any evidence that JNK was directly involved in the inhibition of cell migration observed with UA. The contrary appears to be the case, rather than suppression of JNK by UA as reported by Yeh and colleagues, our data indicates that JNK activity is promoted by UA in GBM cells and mediates cytotoxicity. No adverse interaction between UA and the JNK inhibitor SP600125 was observed, demonstrating the potent inhibition of migration activated by UA, while also suggesting that UA likely affects GBM migration using a JNK-independent signalling pathway in GBM cells. Moreover, subtoxic doses of UA were very potent inhibitors of GBM cell migration, whereas TMZ, Gefitinib or BCNU were unable to significantly reduce cellular migration.

To conclude, this study has demonstrated that UA stimulates the accumulation of lysosomes which are thought to play a role in UA-induced cell death. This study also demonstrates that UA has a greater capacity to inhibit GBM migration over currently used chemotherapeutic agents and provides rationale for further investigation as a potential therapeutic target.

## Supporting information

Supplementary Movie - Scratch Closure

## Abbreviations

UA: Ursolic acid
GBM: Glioblastoma Multiforme
TMZ: temozolomide
3-MA: 3-methyladenine.

## Acknowledgements

This work is supported by Irish Research Council IRCSET grant (G.E.C), Spanish Ministry of Economy and Competitiveness and European Regional Development Fund Grant number SAF2015-64123-P (C.B. and G.P.C). The authors also thank the FOCAS Research Institute, TU Dublin and Institut de Neurociències at UAB for the use of facilities.

